# Reducing Neutrophil Sialic Acid Residues Alleviates Cerebral Hypoperfusion in Alzheimer’s Models

**DOI:** 10.64898/2026.09.23.753660

**Authors:** Nairuti Nikhil Bhatt, Zeynab Tabrizi, Jolie Janulis, Supriya Chakraborty, Madeleine Weick, Sofia Andrea Franciosa, Gabriela Rodriguez Moore, Christian Agatemor, James E Galvin, Oliver Bracko

## Abstract

**Objective:** Dysregulation of the immune system is increasingly recognized as a contributor to Alzheimers disease (AD) progression, partly through neutrophil adhesion to the cerebral vasculature, which promotes hypoperfusion in AD. Because sialic acid (SA) residues on membrane glycoproteins regulate neutrophil–endothelial interactions, we investigated whether neutrophil sialylation is altered in AD and whether reducing it improves cerebral vascular function.

**Approach and Results:** Lectin blots of isolated neutrophils showed increased SA levels in two AD mouse models, 5xFAD and APP-SAA. We identified α2,3 sialyltransferase-IN-1 as a small-molecule inhibitor that reduces sialylation in vivo. Treating 5xFAD mice with this inhibitor decreased SA on neutrophil membranes, increased cerebral blood flow, and reduced capillary stalling. Leukocytes from patients with preclinical AD and mild cognitive impairment also had higher SA levels than those from age-matched healthy controls.

**Conclusions:** Elevated terminal sialylation of neutrophil glycoproteins contributes to capillary stalling and cerebral hypoperfusion in AD. Neutrophil sialylation may serve as both a biomarker and a therapeutic target for improving cerebral blood flow and slowing disease progression.

## Introduction

Alzheimer’s disease (AD) is a progressive neurodegenerative disorder, which is driven by the accumulation of amyloid-β (Aβ) plaques and neurofibrillary tangles. In recent years, it has become evident that immune cell and vascular dysfunction are not only downstream effects, but critical, interacting contributors to the pathogenesis of aging and neurodegenerative diseases, including AD (1, 2). Vascular contributions to cognitive impairment and dementia (VCID) are thought to precede AD-related pathology, presenting as cerebral hypoperfusion and increased vascular inflammation (3, 4).

Glycosylation residues are one of the most common posttranslational modifications in all cells (5). Emerging evidence indicates that pathological changes in glycoconjugates, such as glycoproteins and glycolipids are strongly related to AD initiation and progression (6). Glycosylation of neutrophils and their functions is multifactorial, ranging from cell-cell communication and cell adhesion, and as such, is critical for neutrophil function and immunity (7-9). Uniformly, glycosylation changes in AD have been linked to facilitate many crucial cellular mechanisms and contribute to pathology. However, abnormal glycosylation has been associated with regulating immune cell activation and inducing inflammatory responses in AD (10).

Sialic acids (SAs) are terminal glycan residues that cap cell-surface glycoproteins and glycolipids, modulating receptor accessibility and mediating cell–cell signaling (Figure 1A) (11). Neutrophils, the most abundant circulating innate immune cell, rely heavily on these glycan-mediated interactions to regulate vascular adhesion, endothelial trafficking, and inflammatory activation within the cerebral microvasculature (12). We have previously shown that neutrophils adhere to and stall cortical capillaries, reducing cerebral blood flow (CBF) and driving neuroinflammation and cognitive decline in AD mouse models (13-15). Although neutrophil– endothelial interactions represent a potential therapeutic target, the molecular mechanisms underlying altered neutrophil adhesion remain poorly understood, and SA dynamics in neutrophils are largely uncharacterized in the context of AD pathophysiology. Here, we hypothesize that altered SA expression on neutrophils promotes aberrant adhesion and contributes to cerebral hypoperfusion in AD (Figure 1A) (16).

**Figure 1.**
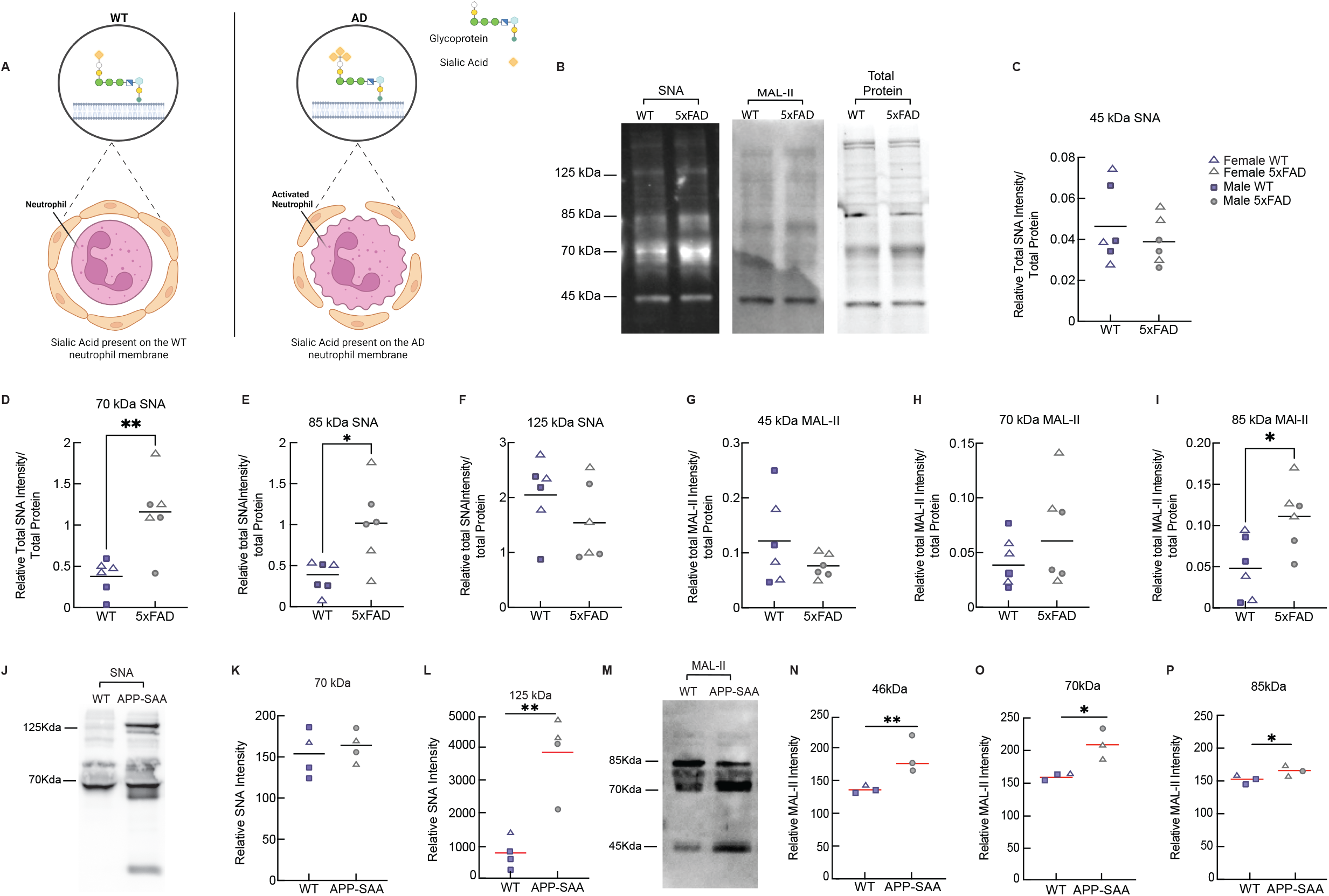
Neutrophil Sialylation and Cerebral Blood Flow in Alzheimers Models. A, Schematic representation of sialic acid (SA) dynamics in neutrophils from 5xFAD mice compared with wild-type (WT) controls. B, SNA (left), MAL-II (middle), and stain-free total protein gel (right) lectin blot of isolated neutrophil membrane fractions from 5-to 6-month-old 5xFAD vs WT mice (n = 6). C, Quantification of relative SNA signal intensity at 45 kDa. D, 70 kDa SNA, E, 85 kDa SNA, F, 125 kDa SNA, G, 45 kDa MAL-II, H, 70 kDa MAL-II, and I, 85 kDa MAL-II. J, SNA lectin blot of isolated neutrophil membrane fractions from APP-SAA vs WT mice (n = 4). K, Quantification of relative SNA signal intensity at 70 kDa, and L, 125 kDa. M, MAL-II lectin blot of isolated neutrophil membrane fractions from APP-SAA vs WT mice (n = 3). N, Quantification of relative MAL-II signal intensity at 46 kDa, O, 70 kDa, and P, 85 kDa.

## Materials and Methods

Materials and Methods are available in the online-only Data Supplement.

## Results

### α-2,6- and α-2,3-linked sialic acid are increased on neutrophil membranes in 5xFAD and APP-SAA mice

We performed lectin blots to measure SA levels on the isolated membrane fractions of neutrophils from 5xFAD and wild-type (WT) mice using Sambucus Nigra Agglutinin lectin (SNA), which preferentially binds SA in α-2,6 linkages to galactose or N-acetylgalactosamines of glycoproteins, and Maackia amurensis Lectin II (MAL-II), which prefers SA in α-2,3 linkages (Figure 1B). SNA-detected SA was significantly increased at 70 kDa and 85 kDa in 5xFAD neutrophil membrane fractions compared to WT control, as confirmed by pixel intensity analysis (Figure 1C-F). MAL-II lectin levels showed increased SA content in 5xFAD membrane fractions, specifically 85 kDa (Figure 1G-I). To test whether increased SA levels are conserved across AD models, we analyzed SNA and Mal-II from 9-month-old APP-SAA mice (knock-in model). Consistent with our previous findings, SA levels were similarly elevated; however, we detected the highest SA content at 125 kDa rather than 70 kDa, as observed in 5xFAD mice (Figure 1J-L). APP-SAA MAL-II also showed overall elevated SA levels compared with controls (Figure 1M-P).

### The α-2,3 sialyltransferase inhibitor IN-1 Reduces neutrophil sialylation and improves cerebral microvascular function in 5xFAD mice, and SA is elevated in human AD

To evaluate whether neutrophil SA content impacts CBF and capillary stalling, the α-2,3 sialyltransferase inhibitor IN-1 (10 mg/kg) was administered to 5–6-month-old 5xFAD and WT mice. Our testing of the compound on isolated neutrophil membranes showed that at 48 hours post-injection, the maximal MAL-II level reduction was observed (Figure 2A, B). To assess the effect on CBF, 10 mg/kg IN-1 or saline was injected 48 hours before cranial window implantation in 5-to 6-month-old 5xFAD and WT mice. Texas Red dextran (70 kDa, plasma) and Rhodamine-6G (leukocytes) were retro-orbitally injected to visualize the microvasculature. Using in vivo two-photon microscopy of lightly anesthetized mice, three-dimensional volumetric stacks and line scans were acquired above the somatosensory cortex (100–400 µm depth) to measure capillary CBF and stalling (Figure 2C, D). IN-1-injected 5xFAD mice showed a trend toward fewer capillary stalls (P = 0.056) compared to saline-injected 5xFAD controls, while WT controls were unaffected (Figure 2E). Line-scan analysis (Figure 2F), to determine CBF, revealed a significant increase in capillary speed and volumetric blood flow in IN-1-injected 5xFAD mice compared to saline controls (Figure 2G, H). Capillary diameter did not differ significantly across groups (Figure 2I). To confirm that IN-1s effects are caused by SA reduction, we performed lectin blots for a second compound, 3Fax-Peracetyl Neu5Ac (12.5 mg/kg, i.p.), in 5–6-month-old 5xFAD mice. The result confirmed that sialyltransferase inhibition reduces SA content, as shown by MAL-II lectin blots (Figure 2J).

**Figure 2.**
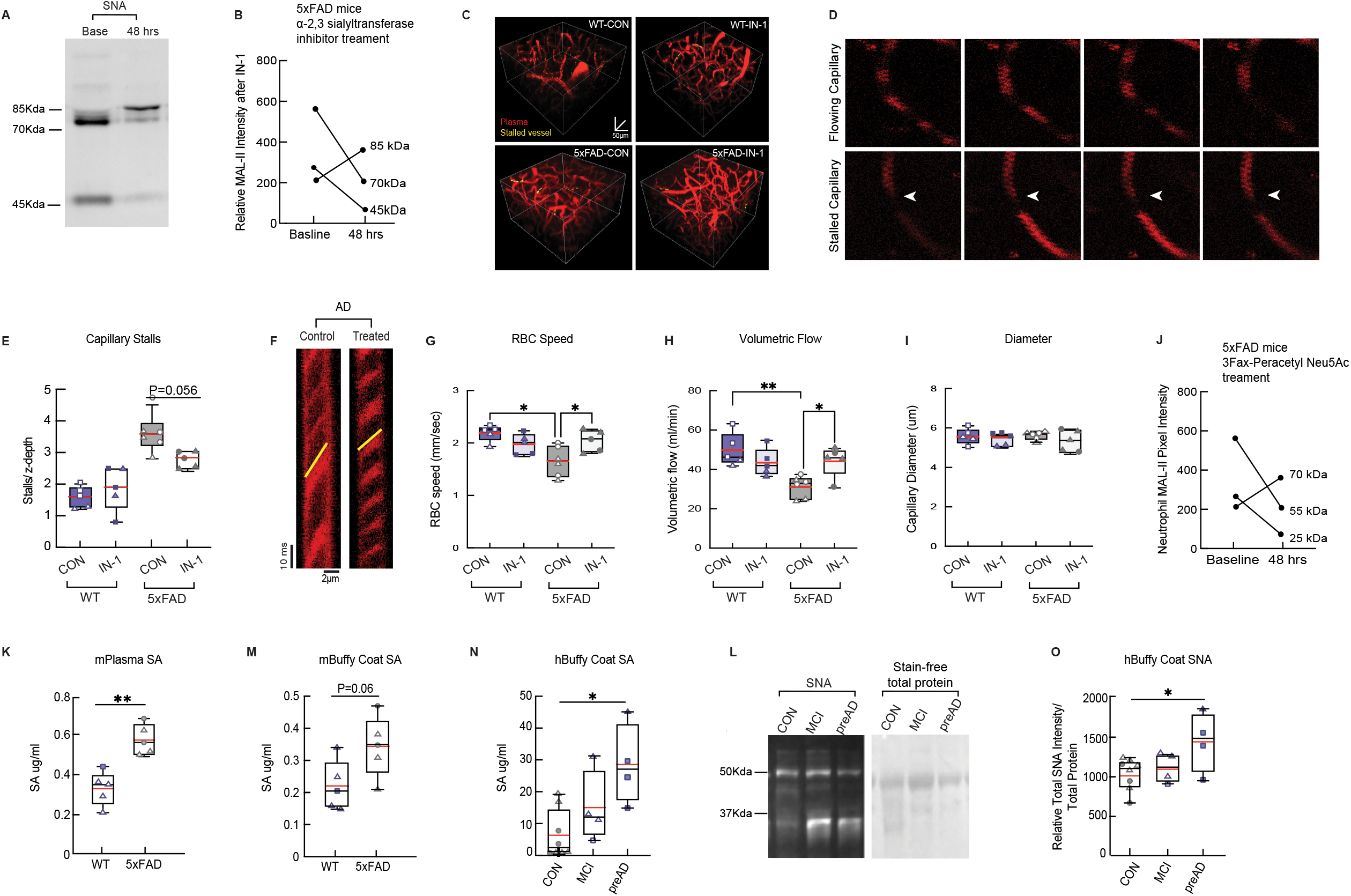
Reducing Sialic Acid Systemically Improves CBF. A, MAL-II lectin blot of isolated neutrophil membrane fractions from 5xFAD at baseline and 48 hrs after IN-1 injection. B, Quantification of the relative MAL-II signal intensity at baseline and after IN-1 injection. Students t-test. C, 3D rendering of in vivo two-photon microscopy imaging stacks from cortical vasculature of 5–6-month-old 5xFAD vs WT mice, imaged 48 hours following intravenous administration of IN-1 (10 mg/kg) or saline. Stalled capillaries are highlighted in yellow. D, Representative time-series images of flowing and stalled capillaries. E, The fraction of stalled capillaries was quantified across groups. Arrowheads point to stalled capillaries. F, Representative line scan images used to determine capillary RBC velocity from 5xFAD and WT mice (scale bars: 2 µm, 10 ms), G, Capillary RBC speed, H, Capillary volumetric blood flow, and I, diameter (5xFAD-CON (n = 5; 125 vessels); 5xFAD-IN-1 (n = 5; 141 vessels); WT-CON (n = 6; 139 vessels); WT-IN-1 (n = 5; 127 vessels) mice. For 2 PM data, two-way ANOVA with multiple comparisons (Genotype; Treatment). Significance was determined as ^*^P < 0.05, ^**^P < 0.01; box plot: whiskers extend 1.5× the difference between the 25th and 75th percentiles of the data, the red horizontal line represents the median, and the black line represents the mean. J, Quantification of the relative SA signal intensity at baseline and 48 hours after 3Fax-Peracetyl Neu5Ac injection. K, ELISA quantification of plasma SA and L, SA content in buffy coat from 5xFAD and WT mice. n = 6; Students t-test; ^**^P < 0.01. M, SA content in human buffy coat measured by ELISA, N, Example SNA lectin blot with stain-free total protein image, O, quantification of SNA levels from control (n = 7), MCI (n = 4), and preclinical AD (n = 4) patients. One-way ANOVA with multiple comparisons; ^**^P < 0.01.

SA ELISA on plasma and buffy coats from 5xFAD and WT mice revealed significantly increased SA in plasma and a trend toward elevated levels in buffy coats (Figure 2K, L). In human samples, SA ELISA and lectin blots from frozen buffy coats showed a trend toward increased SA in mild cognitive impairment (MCI) and a significant increase in preclinical AD patients compared to healthy controls (Figure 2M-O).

## Discussion

Neutrophils undergo dynamic regulation of sialylation during granule formation and maturation, and these changes in sialylation status have important downstream consequences for protein stability, intracellular trafficking, and immune function (17-19). Given these findings, aberrant sialylation has emerged as a potentially critical but underexplored factor in disease contexts, particularly in neurodegenerative conditions such as Alzheimers disease (AD) (16, 20, 21). Here, we have shown that neutrophils from 5xFAD and APP-SAA mouse models exhibit increased sialic acid (SA) content, indicating altered glycosylation patterns that have not been previously characterized in AD mouse models. To investigate the physiological impact of this elevated SA on cerebral blood flow (CBF), we employed both in vivo and translational approaches, finding that enzymatic reduction of SA residues in vivo leads to increases in CBF. Importantly, this observation is supported by clinical relevance, as elevated SA levels were also detected in buffy coats from patients with preclinical AD. Together, these findings suggest that aberrant sialylation may represent a conserved mechanism contributing to cerebrovascular dysfunction in AD.

Sialylation has also been implicated in microglial responses in AD, and a recent study identified increased SA on microglia in an AD mouse model (20, 22). Notably, the study showed that 5xFAD mice had increased α-2,6 sialic acid specifically on plaque-associated microglia, compared with WT mice. Therefore, this excess sialylation may inhibit normal microglial immune recognition of amyloid plaques in AD (20, 21). We have demonstrated for the first time that neutrophils from 5xFAD and APP-SAA mice similarly showed increased SA, and desialylation led to increased CBF. We hypothesize that this elevated SA on neutrophils alters their function, such as cell adhesion, invasion, and transmigration.

A recent publication identified hyperglycosylation as an emerging pattern in AD mouse models and patients (23), consistent with our findings of elevated SA on neutrophils from 5xFAD and APP-SAA mice. We hypothesize that this increased sialylation alters neutrophil adhesion, supported by evidence that ST3Gal-IV-deficient neutrophils show significantly reduced P- and E-selectin binding. ST3Gal-IV is a key α-2,3-sialyltransferase responsible for sialylation of selectin ligands on glycoproteins and glycolipids, demonstrating that ST3Gal-IV-mediated sialylation directly contributes to neutrophil adhesion (24). Consistent with this, IN-1-mediated desialylation reduced neutrophil SA and improved CBF, likely through detachment of adherent neutrophils from capillary walls.

We have shown that SA increases in neutrophils and correlates with capillary stalling in cerebral capillaries. We have further determined that IN-1 injections administered 48 hours before imaging can improve CBF. However, one shortcoming of our approach is that IN-1, because it was injected intravenously, desialylates not only neutrophils but also other blood cells, pericytes, and endothelial cells. Endothelial cells form the glycocalyx, a heavily sialylated structure. Because SA regulates cell-cell interactions, further investigation is needed to determine whether IN-1 alters its homeostatic function (25, 26). While neutrophils and other leukocytes are likely not the sole contributors to changes in CBF, we showed that leukocytes mostly contribute to stalled capillaries regardless of treatment, suggesting that neutrophils play a critical role. Increased SA levels in buffy coats from mice and humans confirm the results from isolated neutrophils. The buffy coat contains not only leukocytes but also platelets; however, neutrophils make up the largest fraction of leukocytes. However, this approach cannot rule out the possibility that other cells may be responsible for the increased SA levels in the patient samples. In summary, we have demonstrated that SA on neutrophils is increased in AD mouse models and patients with MCI and preclinical AD and contributes to vascular damage in cortical capillaries and reductions in CBF.

Elevated SA in buffy coats from both mice and preclinical AD patients further supports these findings. If confirmed, the results suggest that SA measures on neutrophils could serve as an early blood biomarker.

## Significance

Capillary stalling by neutrophils reduces cerebral blood flow in Alzheimers disease (AD) (14, 27), but why neutrophils adhere so readily has been unclear. This study shows that neutrophils from two AD mouse models (5xFAD and APP-SAA) carry more terminal sialic acid on their membrane glycoproteins. Blocking α2,3 sialylation with a small-molecule inhibitor lowered neutrophil sialic acid and improved capillary blood flow in 5xFAD mice. Immune cells from the buffy coats from patients with preclinical AD also showed elevated sialic acid. These findings identify neutrophil sialylation as a candidate mechanism of cerebral hypoperfusion in AD, and as a possible early blood biomarker and therapeutic target.

## Acknowledgments

We thank all members of the BloodFlowLab for scientific discussions and feedback on the manuscript.

## Author contributions

Nairuti Nikhil Bhatt (Conceptualization; Data curation; Formal analysis; Writing – review & editing); Zeynab Tabrizi (Conceptualization; Data curation; Formal analysis; Writing – original draft); Supriya Chakraborty (Data curation; Formal analysis; Writing – review & editing); Jolie Janulis (Data curation; Formal analysis; Writing – review & editing); Madeleine Weick (Data curation; Writing – review & editing); Sofia Andrea Franciosa (Data curation; Writing – review & editing); Gabriela Rodriguez Moore (Data curation; Writing – review & editing); Christian Agatemor (Conceptualization; Supervision; review and editing); James E Galvin (Data curation; review and editing); Oliver Bracko (Conceptualization; Project administration, Supervision, and Validation; Writing – original draft).

## Data availability statement

The data that support the findings of this study are available from the corresponding author upon reasonable request.

## References

1. Van Eldik LJ, Carrillo MC, Cole PE, Feuerbach D, Greenberg BD, Hendrix JA, et al. The roles of inflammation and immune mechanisms in Alzheimers disease. Alzheimers Dement (N Y). 2016;2(2):99–109.

2. Jorfi M, Maaser-Hecker A, Tanzi RE. The neuroimmune axis of Alzheimers disease. Genome Med. 2023;15(1):6.

3. Anwer M, Poliakova T, Albanus RD, Bhuiyan MIH, Brickman AM, Chaudhuri S, et al. Vascular contribution to cognitive impairment and dementia (VCID): proceedings of 2025 workshop of the Jackson Laboratory. Mamm Genome. 2026;37(1).

4. Santisteban MM, Iadecola C. The pathobiology of neurovascular aging. Neuron. 2025;113(1):49–70.

5. He M, Zhou X, Wang X. Glycosylation: mechanisms, biological functions and clinical implications. Signal Transduct Target Ther. 2024;9(1):194.

6. Singh S, Singh SK. The Glyco-Remodeling Hypothesis in Alzheimers Disease: Linking Protein N-Glycosylation to Pathogenesis and Translation. ACS Chemical Neuroscience. 2026;17(18):3252–71.

7. Villanueva-Cabello TM, Gutiérrez-Valenzuela LD, Salinas-Marín R, López-Guerrero DV, Martínez-Duncker I. Polysialic Acid in the Immune System. Front Immunol. 2021;12:823637.

8. Sakarya S, Rifat S, Zhou J, Bannerman DD, Stamatos NM, Cross AS, et al. Mobilization of neutrophil sialidase activity desialylates the pulmonary vascular endothelial surface and increases resting neutrophil adhesion to and migration across the endothelium. Glycobiology. 2004;14(6):481–94.

9. Choi H, Kim C, Song H, Cha MY, Cho HJ, Son SM, et al. Amyloid β-induced elevation of O-GlcNAcylated c-Fos promotes neuronal cell death. Aging Cell. 2019;18(1):e12872.

10. Cheng S, Xiao B, Luo Z. Glycosylation in neuroinflammation: mechanisms, implications, and therapeutic strategies for neurodegenerative diseases. Transl Neurodegener. 2025;14(1):47.

11. Botella Lucena P, Heneka MT. Inflammatory aspects of Alzheimers disease. Acta Neuropathol. 2024;148(1):31.

12. Lightfoot A, McGettrick HM, Iqbal AJ. Vascular Endothelial Galectins in Leukocyte Trafficking. Front Immunol. 2021;12:687711.

13. Ruiz-Uribe NE, Bracko O, Swallow M, Omurzakov A, Dash S, Uchida H, et al. Vascular oxidative stress causes neutrophil arrest in brain capillaries, leading to decreased cerebral blood flow and contributing to memory impairment in a mouse model of Alzheimers disease. bioRxiv. 2023.

14. Cruz Hernandez JC, Bracko O, Kersbergen CJ, Muse V, Haft-Javaherian M, Berg M, et al. Neutrophil adhesion in brain capillaries reduces cortical blood flow and impairs memory function in Alzheimers disease mouse models. Nat Neurosci. 2019;22(3):413–20.

15. Tabrizi Z, Lim XR, Chakraborty S, Harraz OF, Bracko O. Systemic Piezo1 activation improves cerebrovascular function in Alzheimers disease. Alzheimers Dement. 2025;21(12):e71016.

16. Zhu W, Zhou Y, Guo L, Feng S. Biological function of sialic acid and sialylation in human health and disease. Cell Death Discov. 2024;10(1):415.

17. Chakraborty S, Tabrizi Z, Bhatt NN, Franciosa SA, Bracko O. A Brief Overview of Neutrophils in Neurological Diseases. Biomolecules. 2023;13(5).

18. Zhang F, Xia Y, Su J, Quan F, Zhou H, Li Q, et al. Neutrophil diversity and function in health and disease. Signal Transduct Target Ther. 2024;9(1):343.

19. Butovsky O, Rosenzweig N, Kleemann KL, Jorfi M, Kuchroo VK, Tanzi RE, et al. Immune dysfunction in Alzheimer disease. Nat Rev Neurosci. 2026;27(3):196–218.

20. Fastenau C, Wickline JL, Smith S, Odfalk KF, Solano L, Bieniek KF, et al. Increased alpha-2,6 sialic acid on microglia in amyloid pathology is resistant to oseltamivir. Geroscience. 2023;45(3):1539–55.

21. Fastenau C, Crisp R, Keating M, Ochoa E, Richardson TE, Flanagan ME, et al. Sialylation patterns in cerebral amyloid angiopathy. Brain Pathol. 2026;36(2):e70042.

22. Puigdellívol M, Allendorf DH, Brown GC. Sialylation and Galectin-3 in Microglia-Mediated Neuroinflammation and Neurodegeneration. Front Cell Neurosci. 2020;14:162.

23. Hawkinson TR, Liu Z, Ribas RA, Medina T, Nielsen RS, Clarke HA, et al. Hyperglycosylation is a metabolic driver of Alzheimers disease. Nat Metab. 2026;8(6):1410–25.

24. Ellies LG, Sperandio M, Underhill GH, Yousif J, Smith M, Priatel JJ, et al. Sialyltransferase specificity in selectin ligand formation. Blood. 2002;100(10):3618–25.

25. DAddio M, Frey J, Otto VI. The manifold roles of sialic acid for the biological functions of endothelial glycoproteins. Glycobiology. 2020;30(8):490–9.

26. DAddio M, Frey J, Tacconi C, Commerford CD, Halin C, Detmar M, et al. Sialoglycans on lymphatic endothelial cells augment interactions with Siglec-1 (CD169) of lymph node macrophages. FASEB J. 2021;35(11):e22017.

27. Bracko O, Cruz Hernandez JC, Park L, Nishimura N, Schaffer CB. Causes and consequences of baseline cerebral blood flow reductions in Alzheimers disease. J Cereb Blood Flow Metab. 2021;41(7):1501–16.

