## SUPPLEMENTARY MATERIAL for "Reducing Neutrophil Sialic Acid Residues Alleviates Cerebral Hypoperfusion in Alzheimer’s Models"

**Supplemental Materials and Methods**

***Animals***

C57BL/6J 5xFAD and wild-type mice were purchased from Jackson Laboratory (Bar Harbor, ME), and the colony was maintained at the University of Miami. The mice were used at 5-6 months of age for all experiments. We used 6-month-old APP-SAA mice purchased from Jackson Laboratory as a second model. All animal procedures were in accordance with the National Institutes of Health Guide for the Care and Use of Laboratory Animals and approved by the Institutional Animal Care and Use Committee (IACUC) at the University of Miami and the University of Southern California.

***Neutrophil Isolation and Membrane Extraction***

The whole blood (approximately 100-120 μl) was extracted from 5xFAD mice ,APP-SAA mice and their controls using a submandibular blood draw and collected in 1x EDTA tubes (approximately 30 μl) to prevent blood coagulation. Ice-cold RBC lysis buffer was added to the whole blood at a 1:9 ratio, where every 1 ml of blood had 9 ml of lysis buffer. The solution was incubated for 10 minutes at room temperature, and 1xPBS was added to stop the RBC lysis reaction. Next, the lysed blood was centrifuged at 1100 rpm for 20 minutes at 4 °C. The supernatant was discarded, and the buffy coats were carefully extracted from the pellet, which resuspended slowly in a 2 ml EasySep™ (STEMCELL) buffer. The neutrophils were then extracted using the Easy Sep protocol's negative selection. Next, the cell viability was tested by mixing the cells in a 1:1 ratio with trypan blue (Bio-Rad) and counting with a phase-contrast microscope (ZEISS). The membranes was extracted following a manufacturer's protocol (Thermo Fischer). The neutrophil membrane proteins were stored at -20 °C.

***Mouse Sialic Acid Lectin Blot***

Immediately post-isolation, neutrophil pellets were lysed in 50 µL of RIPA with 1x concentration of Thermo Scientific™ Pierce Protease Inhibitor Mini Tablets and aliquoted into two 25 µL volumes prior to incubation and freezing. Upon thawing, proteins were normalized using the Pierce BCA Protein Assay Kit (Thermo Scientific™) following a modified manufacturer's protocol to account for low sample availability (To ensure accurate protein normalization to blot intensity, blots were later quantified using a Bio-

Rad StainFree blot read via ChemiDoc). The membrane protein samples were first heated at 95 °C for 5 minutes, and the lectin blot was then run at 100V on a 12% or 4–20% TGX Bio-Rad StainFree Gel for up to 90 minutes. Transfer was conducted for 10 minutes using the Bio-Rad Turbo Transfer System to achieve optimal transfer across all molecular weights. After transfer, the PVDF membrane was blocked for 1 hour at room temperature in a 3% BSA blocking solution with 1x TBST. To detect sialic acid, membranes were incubated with 12 µL/mL SNA-EBL, fluorescein (Vector Labs, 2 mg/mL) or 12 µL/mL MAL-II biotinylated lectin (Vector Labs, 1 mg/mL) for 1 hour at room temperature with gentle shaking. Blots were first imaged on the ChemiDoc Fluorescein setting to detect SNA lectin binding, prior to incubation with a 1:1000 dilution of HRP-conjugated streptavidin for one hour at room temperature. Biotinylated Mal-II signal was detected using Thermo Scientific™ SuperSignal™ West Pico PLUS Chemiluminescent Substrate, following the manufacturer's protocol. Membranes were developed using the ChemiDoc Imaging System (Bio-Rad) at varying exposure times under 60 seconds, depending on saturation levels. Membranes were analyzed using Image Lab software to calculate pixel density, normalized to total lane protein signal (Bio-Rad StainFree).

***Mouse Sialic Acid ELISA***

200 µL of whole blood was drawn from 5-6 months old five 5xFAD and wild-type mice. Blood was diluted 1:1 in PBS with 0.1% BSA and 0.6% 2 mM EDTA (without $Ca^{2+}$ and $Mg^{2+}$) and centrifuged at 800 x g for 10 minutes at room temperature. The top plasma layer was removed and stored in tubes, and the buffy coat was carefully removed and washed 3 times in PBS with 0.1% BSA. Plasma and buffy coats were both lysed in RIPA with protein inhibitor (Complete, Roche). Samples were sonicated (only buffy coats), vortexed, and then incubated on ice for 15 minutes. Samples were centrifuged at 14,000 g for 20 minutes at 4 °C. The supernatant was transferred into new tubes, and Pierce BCA Protein Assay (Thermo Fisher Scientific) was performed to determine protein concentrations, which were then stored at −80 °C. These plasma and buffy coats were analyzed by sandwich ELISA for Sialic acid (MBS264997; MyBioSource) following the manufacturer’s protocol. The Sialic acid concentration was calculated by comparing the sample absorbance with that of known concentrations of a Sialic acid standard. OD was

measured at 450 nm using a plate reader, and the data were analyzed using Excel (Microsoft) and Prism (GraphPad).

***Sialic Acid Inhibitor Treatments***

*α 2,3 Sialyltransferase IN-1:* We used the SA inhibitor drug " α 2,3 Sialyltransferase IN-1 " (MedChemExpress) at a concentration of 10 mg/kg and injected the drug retro-orbitally at 12, 24, 48, and 72 hours under 3% isoflurane anesthesia. Immediately after the end of each time point, we extracted the blood as mentioned above and subsequently isolated the neutrophils and their membrane proteins to further conduct the lectin blots.

*" 3 Fax-Peracetyl 5 NeuAc ", the Sialyltransferase inhibitor:* We used SA inhibitor 3 Fax-Peracetyl 5 NeuAc (Millipore Sigma) at a final concentration of 12.5 mg/kg mixed with PEG-300 (Sigma Aldrich) in 2-fold, then injected it in the mice intraperitoneally, at 24, 48, 72, and 96 hours under 3% isoflurane anesthesia, one injection per day for each time point. We extracted the whole blood immediately after each time point, as mentioned in the protocol above. Subsequently, we isolated the neutrophils and their membrane proteins to further conduct the lectin blots.

***Cranial Window Implantation***

5xFAD and wild-type mice were injected 48 hours before craniotomy with 10 mg/kg of the drug, " α2,3 Sialyltransferase IN-1 ", or saline as a control, and the drug was administered retro-orbitally. Then, the mice were anesthetized using a Low-Flow Electronic Vaporizer (Kent Scientific) with 1.5% isoflurane while the mice were secured in a custom-built stereotactic surgery frame. The mice received a subcutaneous injection of 0.005 mg/100 g atropine (Sparhawk Laboratories) to reduce lung secretions, 0.025 mg/100 g dexamethasone (Aspen) to control post-surgical inflammation, and 0.5 mg/100 g ketoprofen (Bimeda) for pain relief. A local nerve block was also administered at the incision site with 0.1 ml of 0.125% bupivacaine (Hospira Inc.). The body temperature was maintained at 37°C using a thermometer and a feedback-controlled heating blanket (Kent Scientific). The head was shaved and cleaned three times with alternating solutions of 70% ethanol and iodine (Millipore). A precise 6-mm diameter craniotomy was performed over the cerebral cortex using a high-speed drill (HP4–917-21, Fordom). A sterile 8-mm diameter glass coverslip (Electron Microscopy Sciences) was placed and secured onto the skull using tissue adhesive (3M Vetbond) and dental cement (Co-Oral-Ite Dental). All steps were performed with sterile techniques. After the craniotomy, the mice were taken to the microscopy room.

***In Vivo Two-photon Microscopy***

Mouse anesthesia was maintained at 1.2-1.5% during *in vivo* two-photon microscopy. Body temperature was maintained at 37°C using a feedback-controlled heating pad (Kent Scientific). To visualize the microvasculature, a retro-orbital injection of Texas Red dextran (50 μl, 2%, MW = 70,000 kDa, Thermo Fisher Scientific) in saline was administered before imaging. Additionally, leukocytes and blood platelets were labeled *via* a retro-orbital injection of Rhodamine 6G (0.1 ml, 1 mg/ml in 0.9% saline, Millipore Sigma), with leukocytes being distinguished from blood platelets using a retro-orbital injection of Hoechst 33342 (50 μl, 4.8 mg/ml in 0.9% saline, Thermo Fisher Scientific).

Three-dimensional images of the cortical vasculature and measurements of red blood cell flow speeds in specific vessels were acquired using a two-photon excited fluorescence microscope. Imaging was performed with 830-nm, 120-fs pulses from an InSight®X3+ laser oscillator (Spectra-Physics). The laser beams were scanned by galvanometric scanners (1 frame/s) and focused on the sample using a 20× water-immersion objective lens (numerical aperture (NA) of 1.0; Carl Zeiss Microscopy). The emitted fluorescence was captured using a four-channel detection system based on a Bergamo II (Thorslabs). A first long-pass dichroic at 562 nm was followed by a second long-pass dichroic at 495 nm and 635 nm. Bandpass emission filters were 447/60, 525/50, 607/70, and 647 LP. Data acquisition was controlled using Thorslabs software. Image stacks were taken at one μm intervals axially to a cortical depth of 200–600 μm to visualize the cortical vasculature.

***Capillary Stalling and Blood Flow Analysis***

We used ImageJ software Fiji (https://fiji.sc/) to process and quantify capillary stalling of the three-dimensional imaging stacks of the cortical vasculature. We manually determined capillaries as stalled or flowing if the stall was at least for five frames in any position within the capillary segment. We counted at least four image stacks for each mouse, and another experiment independently verified all stalled vessels. Capillary diameters were determined using ImageJ's plot profile function. 15-25 capillaries were measured per mouse using line scans along the vessel's center axis to measure RBC flow speeds in capillaries. The movement of RBCs was tracked over successive line scans by constructing a space-time plot, where the x-axis represented distance along the vessel axis, and the y-axis represented time. Moving RBCs formed diagonal stripes on this plot, with the slope inversely proportional to the RBC speed, which was calculated using a radon transform-based algorithm.

***Patient Buffy Coat Protein Isolation***

Stock samples of isolated buffy coat from patients were assessed for cell count and viability by scraping frozen samples. Each scrape was resuspended in 1 mL of 1X PBS with 1X complete protease inhibitors (Roche) and incubated on ice for 30 minutes. 1 µl of that resuspension was added to a diluted trypan blue solution for visual inspection under a dissection microscope. Samples were then divided in half; one half was immediately returned to -80°C, while the other was lysed in 5 volumes of RIPA buffer with complete protease inhibitors (Roche) and incubated on ice for 30 minutes. The lysate was centrifuged for 20 minutes at 1000 × g in a pre-chilled microcentrifuge at 4°C. The supernatant from each sample was collected and diluted 10-fold using 1X PBS with 1X complete protease inhibitors (Roche) and then aliquoted for storage at -80°C. One aliquot of each sample was then assayed to determine protein concentration using the Pierce BCA Assay, which was read and analyzed following the manufacturer's instructions. Individual samples were diluted with DI water to obtain an equal protein concentration.

**Table Patients**

| | | **First Diagnosis** | **Last Diagnosis** | **ApoE** | **Amyloid present** | **Final Dx** |
|---|---|---|---|---|---|---|
| 1 | HBI0001 | Control | Control | 4/4 | Yes | Preclinical AD |
| 2 | HBI0005 | Control | Control | 3/4 | Yes | Preclinical AD |
| 3 | HBI0006 | Control | Control | 3/3 | No | True Control |
| 4 | HBI0007 | Control | Control | 3/3 | No | True Control |
| 5 | HBI0010 | Control | Control | 3/3 | No | True Control |
| 6 | HBI0013 | Control | Control | 3/3 | Yes | Preclinical AD |

| | | | | | | |
|---|---|---|---|---|---|---|
| 7 | HBI0017 | Control | Control | 3/3 | No | True Control |
| 8 | HBI0019 | MCI | MCI | 3/3 | No | MCI |
| 9 | HBI0024 | Control | Control | 3/3 | No | True Control |
| 10 | HBI0034 | MCI | MCI | 3/3 | No | MCI |
| 11 | HBI0046 | MCI | MCI | 3/4 | No | MCI |
| 12 | HBI0053 | MCI | MCI | 3/3 | Yes | MCI |
| 13 | HBI0056 | Control | Control | 3/4 | No | True Control |
| 14 | HBI0060 | Control | Control | 3/4 | Yes | Preclinical AD |
| 15 | HBI0063 | Control | Control | 3/3 | No | True Control |
| 16 | HBI0069 | Control | Control | 3/3 | No | True Control |

***Patient Sialic Acid Lectin blot***

Individual samples were diluted with DI water containing complete protease inhibitors (Roche) to a final protein concentration of 5 µg/µl. 3.33uL of 1;9 β-mercaptoethanol to 4x Lammeli buffer (BioRad) was added to 10uL of the diluted samples. The lysed samples were loaded onto 12-well Criterion 12% SDS-PAGE gels (Bio-Rad) and resoled at a constant 150 V for 60 min. Proteins were transferred to 0.2 µm nitrocellulose membranes using the Trans-Blot Turbo Transfer System (Bio-Rad). Membranes were blocked in 3% Bovine Serum Albumin (BSA) in Tris-buffered saline with 0.1% Tween-20 (TBS-T) for 1 h at room temperature. The membranes were agitated during incubation in a 3% BSA Blocking Buffer + TBST solution for one hour at room temperature. Membranes were then incubated with 15 µl SNA biotinylated lectin (Vector Labs) or 4.5 µl MAL-II lectin in 10 ml of blocking buffer for one hour at room temperature with gentle shaking. For lectin detection, mixed a 1:10000 dilution of HRP-conjugated streptavidin with 10 ml of 3% BSA Blocking Buffer + TBST for each membrane, then incubate for 40 minutes at room temperature with gentle shaking. Immunoreactive bands were visualized by enhanced chemiluminescence (SuperSignal West Pico PLUS), and digital images were acquired using a ChemiDoc Imaging System (Bio-Rad). Samples were analyzed by densitometry in ImageJ.

***Patient Sialic Acid ELISA Assay***

The buffy coat was analyzed by sandwich ELISA for Sialic acid (AB282923; Abcam) following the manufacturer's protocol. The Sialic Acid concentration was calculated by comparing the sample absorbance with that of known concentrations of a Sialic Acid standard. OD was measured at 450 nm using a plate reader.

***Statistical Analyses***

Boxplots were created using Prism 11.0.1 software (GraphPad). The box plots indicate the interquartile range (IQR), encompassing the data's 25th to 75th percentile. Whiskers extend to 1.5 times the IQR. Values outside the whiskers are considered outliers, and the mean is calculated excluding these outliers (outliers are shown in the graphs). The normality of data was assessed using the D'Agostino-Pearson normality test. Statistical comparisons between groups were made using the student's t-test and one-way and two-way ANOVA with multiple comparisons. P-values less than 0.05 were considered statistically significant. Data analysis was conducted under double-blinded conditions, where the experimenter was unaware of the treatment until the completion of data analysis.
